# Aurora B controls microtubule stability to regulate abscission dynamics in stem cells

**DOI:** 10.1101/2024.03.06.583686

**Authors:** Snježana Kodba, Amber Öztop, Eri van Berkum, Malina K. Iwanski, Wilco Nijenhuis, Lukas C. Kapitein, Agathe Chaigne

## Abstract

Abscission is the last step of cell division leading to the complete separation of the two sister cells and consists of the cutting of a cytoplasmic bridge. Abscission is mediated by the ESCRT membrane remodeling machinery which also triggers the severing of a thick bundle of microtubules that needs to be cleared prior to abscission. Here, we show that rather than being passive actors in abscission, microtubules control abscission speed. Using mouse embryonic stem cells, which transition from slow to fast abscission during exit from naïve pluripotency, we investigate the molecular mechanism for the regulation of abscission dynamics and identify a feedback loop between the activity of Aurora B and microtubule stability. We demonstrate that naïve stem cells maintain high Aurora B activity after cytokinesis. This high Aurora B activity leads to transient microtubule stabilization that delays abscission. In turn, stable microtubules promote the activity of Aurora B. When cells exit naïve pluripotency, a decrease in Wnt signaling leads to a decrease in the activity of Aurora B, less stable microtubules, and a faster abscission. Overall, our data demonstrate that Aurora B-dependent microtubule stability controls abscission dynamics.

## Introduction

Cell division is crucial to multicellular development and involves highly conserved protein complexes used to separate the chromosomes and the cytoplasm of one mother cell into two daughter cells. However, during mammalian development, while cells make fate choices, they can completely rewire their cell division machinery within a few hours. In particular, pluripotent stem cells remain connected with their sister cells through cytoplasmic bridges for a protracted amount of time, while stem cells that have exited naïve pluripotency do not (1). Instead, they rapidly sever this connection in a process called abscission. However, we do not know how abscission duration can be rapidly modulated to allow fate changes during development.

Abscission is a complex process that starts at anaphase. At the center of the spindle, several protein complexes are recruited by the overlapping microtubules of the central spindle, including PRC1 which bundles overlapping microtubules (2–4), centralspindlin which contains in particular the microtubule motor KIF23 (5) and the chromosomal passenger complex (CPC) (6). The CPC contains the kinase Aurora B, as well as INCENP, Borealin and Survivin, and is involved in controlling the proper alignment of the chromosomes before anaphase is triggered. The central spindle signals to the cell cortex which triggers the formation and constriction of an actomyosin ring, leading to the formation of two cells connected by a cytoplasmic bridge. The cytoplasmic bridge contains microtubules and a dense structure called the midbody, which forms at the center of the bridge where the microtubules overlap. The midbody notably contains protein from the central spindle, including the CPC, PRC1 and KIF23 (7– 9). The midbody then serves as a platform for the recruitment of complex machinery that will lead to the severing of the cytoplasmic bridge (10). Briefly, KIF23 recruits Cep55 (11), which in turn recruits Tsg101 and Alix (12, 13). Tsg101 and Alix both interact directly or indirectly with members of the ESCRT complex leading to the formation of filaments of ESCRT-III (14–21). These filaments are remodeled by the AAA-ATPase VPS4 (22) and in turn delocalize from the midbody to the midbody arms, which triggers severing and resealing of the plasma membrane (16, 23, 24).

Importantly, the contents of the cytoplasmic bridge must be removed prior to ESCRT-III mediated abscission (25). Actin remodeling is critical for abscission completion (26– 29). Microtubule removal is thought to be a passive event depending on membrane remodeling. Indeed, in parallel to ESCRT-III mediated scission events, CHMP1B, a member of the ESCRT-III family recruits the microtubule severing protein spastin (30, 31). Spastin then mediates microtubule severing and clearing from the cytoplasmic bridge (24, 30– 33). As such, microtubules are not thought to actively contribute to the timing of abscission and their removal is completely dependent on the dynamics of ESCRT recruitment. Another microtubule severing protein, Katanin, can interact with the ESCRT-III binding domain MIT (Microtubule Interacting and Trafficking) (34) and has been involved in abscission (35). However, recent data suggest that in cancer cells, the timing of microtubule severing is uncorrelated from membrane severing and depends on actin dynamics (26), suggesting that multiple mechanisms could be at play in microtubules severing and abscission.

Abscission dynamics are mediated by the recruitment and repositioning of ESCRT-III filaments from the midbody to the future abscission site on the midbody arms. In cancer cells, the proper localization of ESCRT-III filaments is controlled by the CPC. At the midbody, Aurora B and Borealin can directly phosphorylate the ESCRT-III member CHMP4C (23, 36, 37), which leads to its retention at the midbody or in the cytoplasm and delayed abscission. Aurora B can also indirectly control the phosphorylation of VPS4 through ANCHR (38). Furthermore, Aurora B is important for blocking the abscission of cytoplasmic bridges in the germline (39) and in mouse embryos (40). Aurora B can also phosphorylate the microtubule motor KIF20A (41), which inhibits abscission. Surprisingly, spastin, which is important for severing microtubules and promoting abscission (30–32), has also been shown to be required for abscission arrest in cases of mitotic defect (34), suggesting that microtubules could be a downstream target of Aurora B and an underestimate player of abscission dynamics. However, how microtubules regulate abscission dynamics is not known.

Here, we use mouse embryonic stem cells, which can transition from slow to fast abscission during development, as a model system to investigate a direct role of Aurora B-dependent microtubule stability in controlling abscission. We show that in pluripotent cells, Aurora B activity is locally maintained at high levels at the exit of cell division. Aurora B activity regulates the duration of abscission in stem cells by controlling the stability of microtubules, with high Aurora B activity leading to increased numbers of stable microtubules. In turn, stable microtubules prevent abscission directly and indirectly by enhancing Aurora B activity. Finally, we show that Aurora B activity controls the dynamics of exit from naïve pluripotency and that Wnt signaling participates in Aurora B-dependent slow abscission. Together, our results demonstrate that Aurora B can regulate abscission and cell fate through the direct regulation of microtubules.

## Results

### Aurora B activity decreases slowly during bridge maturation in naïve ESCs

To study the dynamics of abscission, we used mouse embryonic stem cells (ESCs hereafter) as a model system as they undergo a switch from a slow (6 h) to a fast (2-3 h) abscission when they exit naïve pluripotency (1) (Figure 1A). By comparing naïve cells (in 2i-Lif media, naïve ESCs hereafter) and exiting cells (in N2B27 media, exiting ESCs hereafter) we can compare the same cell line undergoing either slow or fast abscission. Before abscission, cytoplasmic bridges are formed and matured, during which actin is removed and the diameter of the bridge narrows (1, 24, 28). To be able to compare bridges of the same age, we synchronized cells before mitotic entry using the CDK1 inhibitor RO-3306, then released the cells for 90 min (see Methods). Cells were then fixed immediately (time point 0 h, which corresponds to bridge formation), 2 h, or 4 h later.

**Fig. 1:**
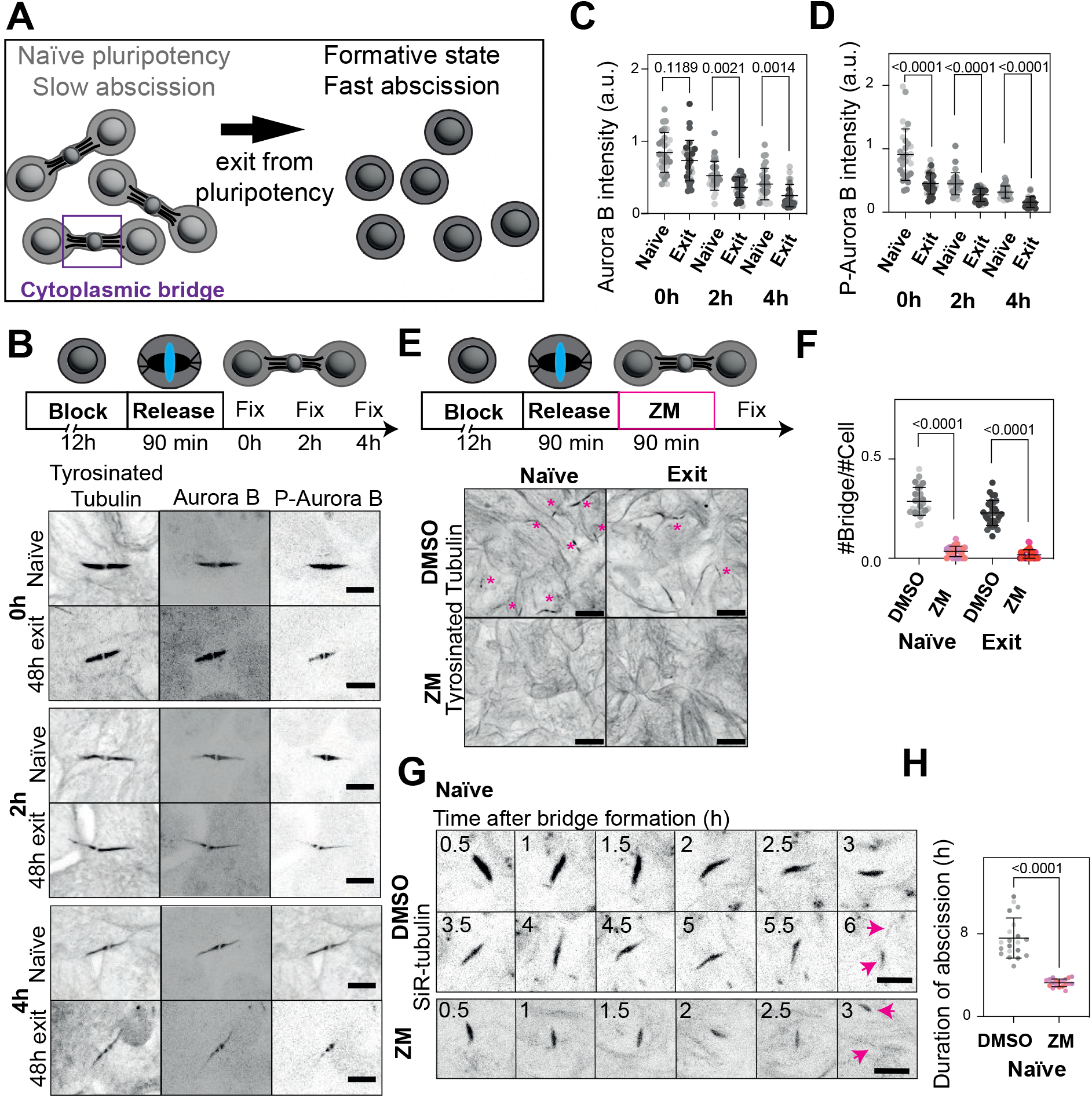
Aurora B activity controls abscission speed. A) Cartoon showing the dynamics of abscission during exit from naïve pluripotency in mouse embryonic stem cells (ESCs). B) Top: schematic of the experimental set-up. Bottom: immunofluorescence showing the localization of Aurora B (middle panel) and P-Aurora B (right panel) in naïve (“naïve”, top) and 48h exiting (“exit”, bottom) ESCs during bridge maturation at bridge formation (0h), 2h after bridge formation (2h) and 4h after bridge formation (4h). The bridge is shown with the staining of tyrosinated tubulin. A Z-projection over the height of the whole cell is shown. Scale bars: 5 *μ*m. C) Quantification of Aurora B intensity in naïve (light grey) and exiting (dark grey) ESCs. The mean and standard deviation are shown. N=3 replicates. D) Quantification of P-Aurora B intensity in naïve (light grey) and exiting (dark grey) ESCs. The mean and standard deviation are shown. N=3 replicates. E) Top: schematic of the experimental set-up. Bottom: immunofluorescence showing the number of bridges in in naïve (left) and 48h exiting ESCs (right) treated with DMSO (top) or 2 *μ*M ZM447439 (bottom). A Z-projection over the height of the whole cell is shown. The bridges are shown with the staining of tyrosinated tubulin. Bridges are highlighted with pink asterisks. Scale bars: 10 *μ*m. F) Quantification of the number of bridges per cell in naïve ESCs treated with DMSO or 2 *μ*M ZM447439 (light grey and light pink, respectively) and exiting ESCs treated with DMSO or 2 *μ*M ZM447439 (dark grey and dark pink, respectively). The mean and standard deviation are shown. N=3 replicates. G) Live-cell imaging of naïve ESCs incubated overnight with 20 nM SiR-tubulin after addition of DMSO (top) or 2 *μ*M ZM447439 (bottom). A Z-projection over the height of the whole cell is shown. Tubulin is shown in black. The pink arrows indicate the cut sites. One frame is shown every 30 min. Scale bars: 10 *μ*m. H) Quantification of the duration of abscission from bridge formation until microtubule severing in naïve ESCs treated with DMSO or 2 *μ*M ZM447439. The mean and standard deviation are shown. N=3 replicates.

Since Aurora B has been implicated in abscission regulation in multiple contexts, we first investigated its involvement in the abscission of ESCs. Aurora B was present over the length of the bridges of naïve ESCs and exiting ESCs (Figure 1B). To measure intensities, we hereafter chose bridges that are flat (contained in a single Z-plane), segmented them in that plane, then measured the mean grey value; thus, all intensities are normalized to the size of bridges. The levels of Aurora B decreased over time in naïve and exiting ESCs, with levels remaining overall higher in naïve cells (Figure 1B,C).

To test whether Aurora B activity was also higher in naïve ESCs, we stained the cells with an antibody that specifically recognizes the phosphorylation of Th232 of Aurora B (42). Similarly to Aurora B, the levels of P-Aurora B decreased over time (Figure 1B,D). Strinkingly, naïve ESCs had higher levels of P-Aurora B than exiting ESCs at all time points of bridge maturation (Figure 1B,D). Finally, we wanted to test whether the higher activity of Aurora B at the bridge of exiting cells was linked to an overall higher activity of Aurora B in exit cells, or whether it was specific to the bridge. To do so, we compared the levels of P-Aurora B in metaphase cells in naïve and exiting cells and found that the levels were similar (Supplementary Figure 1A,B). Thus, our data show that naïve ESC have higher levels of Aurora B activity than exiting ESCs specifically on cytoplasmic bridges, and that this activity decreases during bridge maturation.

### Aurora B regulates abscission dynamics in mouse embryonic stem cells

Next, we tested whether decreasing Aurora B activity impacted abscission duration. Because the CPC is involved in the metaphase-to-anaphase checkpoint, to investigate the role of Aurora B in abscission, we again synchronized cells before mitotic entry using the CDK1 inhibitor RO-3306, then released the cells for 2h before inhibiting Aurora B activity using ZM447439 (hereafter ZM, see Methods). After 90 min of Aurora B inhibition, the localization and amount of Aurora B was unaffected in naïve ESC and slightly decreased in exiting ESCs (Supplementary Figure 1C,E), whereas the activity of Aurora B was strongly reduced both in naïve and exiting ESCs (Supplementary Figure 1D,F). We then counted the number of cytoplasmic bridges as a proxy for abscission duration. The number of bridges was strongly reduced in both naïve and exiting ESCs treated with ZM (Figure 1E,F), suggesting that abscission is faster when Aurora B activity is abolished, in agreement with previous work in other model systems (40, 43). To confirm this, we imaged naïve ESC incubated with the live microtubule probe SiR-tubulin from anaphase onset to microtubule severing with or without ZM. We used the rupture of the microtubule bundle as a proxy for abscission, as membrane rupture always promptly follows the rupture of the microtubule bundle (Supplementary Movie 1, 3/10 cells cut the membrane less than 5 minutes after microtubule rupture and 7/10 cells cut the membrane less than 10 minutes after microtubule rupture, out of N=3 independent experiments). Control cells cut the cytoplasmic bridge in 6h while cells treated with ZM were twice as fast (Figure 1G,H. Movie 1-2). Abscission duration was also shorter in exiting cells when treated with ZM (82 +/-18 min, compared to 171 +/-31 min in control cells) (Movie 3-4).

To confirm the role of Aurora B, we tested whether phosphatases could oppose the effect of Aurora B. To do so, we inhibited the phosphatases PP1 and PP2A using Okadaic Acid (OKA) (44). Naïve ESCs treated with OKA had similar Aurora B intensity at the bridge than controls; exiting ESCs treated with OKA had slightly higher Aurora B intensity than controls (Supplementary Figure 1G,I). In all cases, treatment with OKA led to increased Aurora B activity at the bridge (Supplementary Figure 1H,J). Treatment with OKA also led to an increase in bridge number in exiting ESCs (Supplementary Figure 1K,L), suggesting that inhibiting PP1 and PP2A slows abscission. To confirm the role of phosphatases in abscission dynamics, we added OKA in naïve and exiting cells just after bridge formation and found that treatment with OKA indeed increased the duration of abscission in naïve and exiting cells (Supplementary Figure 1M,N, Supplementary Movie 2-5). Overall, our data show that higher Aurora B activity in pluripotent ESCs delays abscission.

### Wnt-dependent Aurora B activity delays exit from naïve pluripotency

Our data showed that Aurora B activity is higher in naïve ESCs than in ESCs exiting naïve pluripotency. Therefore, we wanted to test whether the activity of Aurora B controls exit from naïve pluripotency. To do so, we performed clonogenicity assays to functionally test the dynamics of exit; briefly, cells were exited from naïve pluripotency for 20h, then replated at clonal density in pluripotency media (2i-Lif) (Figure 2A). Counting the colonies lead to an estimation of the colony forming capacity of the cells, providing a functional test of the dynamics of exit from pluripotency (a faster exit will lead to fewer colonies). To test the role of the activity of Aurora B, we allowed cells to exit from naïve pluripotency for 15h, then treated them with ZM for 90 min, and allowed them to exit during for a further 3h before replating (Figure 2A). This treatment did not affect cell proliferation (Supplementary Figure 2A). When Aurora B activity was decreased by ZM treatment, less colonies formed compared to control (normalized to 1) showing that exit from naïve pluripotency was accelerated (Figure 2B). To confirm these results, we performed qPCRs on several pluripotency related genes (*Rex1, Klf4, Nanog, Esrrb, Klf2, Oct4*, which are genes expressed by naïve cells and *Sox1*, expression of which goes up after exit from naïve pluripotency (45)) after ZM treatment. We found a global downregulation of pluripotency gene expression when Aurora B activity was decreased in naïve and exiting cells (Figure 2B, in all cases the ZM treatment is compared to the DMSO control normalized to 1). Similarly, we noted an upregulation of the expression of Sox1 in exiting cells (bottom row). Conversely, inhibiting PP1 and PP2A using OKA did not affect proliferation (Supplementary Figure 2B) and delayed exit from naïve pluripotency (Figure 2C). Altogether, these results show that Aurora B activity prevents exit from naïve pluripotency.

**Fig. 2:**
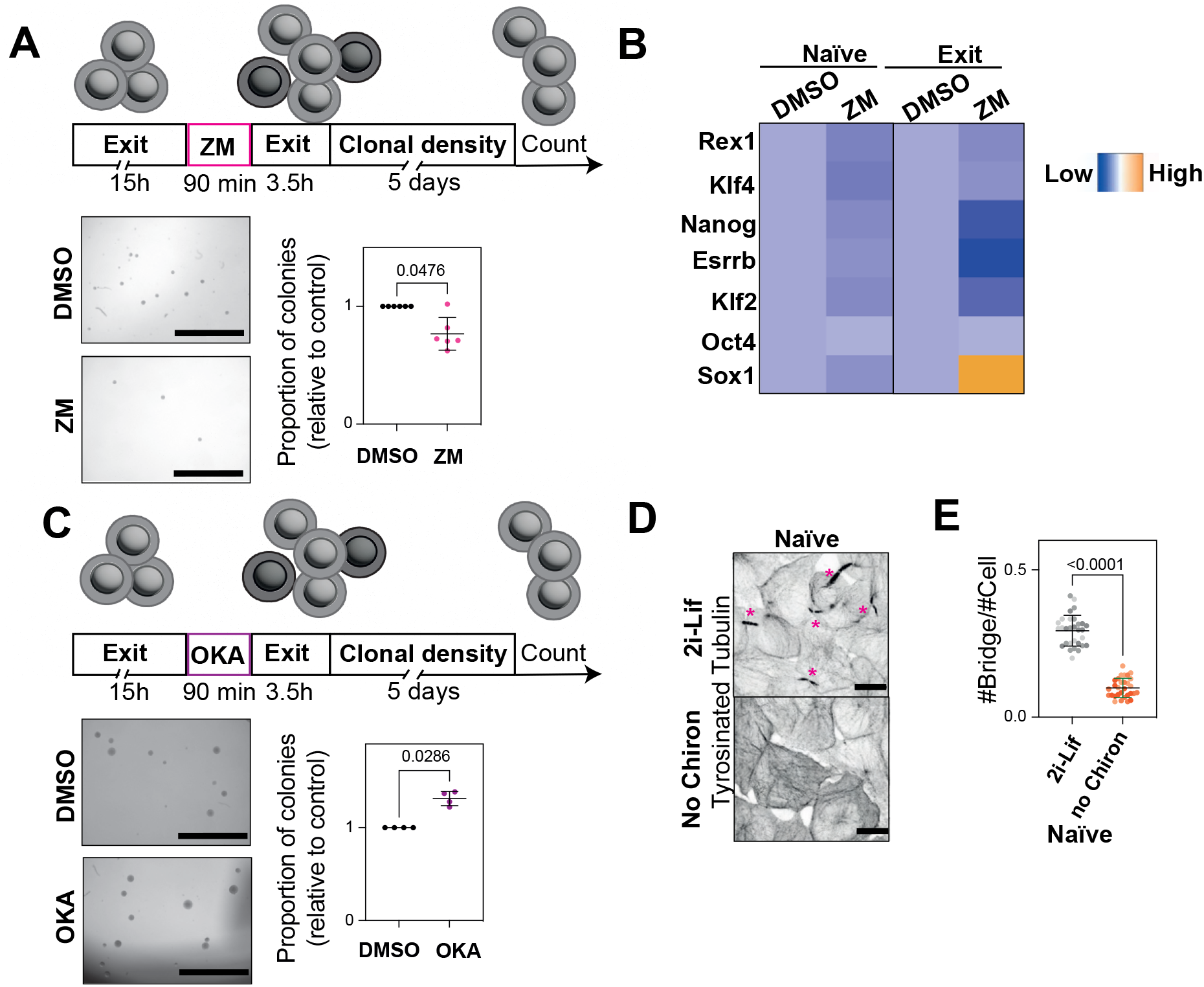
Aurora B activity and Wnt signaling control exit from naïve pluripotency. A) Top: schematic of the experimental set-up. Bottom: clonogenicity assay on ESCs treated for 90 min with 2 *μ*M ZM447439. Left panel: colonies formed by control cells after 20h of exit from naïve pluripotency and cells treated with ZM447439. Scale bars: 1 mm. Right: quantification of the number of colonies formed. N=6. B) Heatmap showing expression levels determined by qPCR of naïve and 48h exiting ESCs treated for 90 min with 2 *μ*M ZM447439. N=3. C) Top: schematic of the experimental set-up. Bottom: clonogenicity assay on ESCs treated for 90 min with 0.5 *μ*M Okadaic Acid (OKA). Left panel: colonies formed by control cells after 20h of exit from naïve pluripotency and cells treated with OKA. Scale bars: 1 mm. Right: quantification of the number of colonies formed. N=4. D) Immunofluorescence showing the number of bridges in naïve ESC in normal 2i-Lif media (top; high Wnt signaling) or without Chiron (bottom; low Wnt signaling). A Z-projection over the height of the whole cell is shown. The bridges are shown with the staining of tyrosinated tubulin. Bridges are highlighted with pink asterisks. Scale bars: 10 *μ*m. E) Quantification of the number of bridges per cell in naïve ESC in normal 2i-Lif media (grey) or after Chiron withdrawal (orange). The mean and standard deviation are shown. N=3 replicates.

Finally, we wondered what could explain the high activity of Aurora B in naïve cells. We reasoned that Wnt signaling could control Aurora B activity; indeed, ESCs are maintained in their naïve state by inhibition of GSK3 (a negative effector of the Wnt pathway) through the inhibitor CHIR-99021 (46–48). Furthermore, Wnt activity has been linked to abscission dynamics in hepatocyte organoids (49). To test this, we decreased Wnt signaling by culturing naïve ESCs without CHIR-99021 (“No Chiron”). We found that removal of CHIR-99021 led to a strong decrease in bridge number (Figure 2D,E), as well as a shortening of the remaining bridges as assessed by the length of the microtubule bundle (Supplementary Figure 2C). Importantly, removal of Chiron did not impair the pluripotency state of the cells nor their ability to proliferate (Supplementary Figure 2D-E). Removing Chiron also led to a decrease in Aurora B and P-Aurora B intensity (Supplementary Figure 2F-H), suggesting that Wnt signaling acts upstream of Aurora B. In conclusion, our findings suggests that the high Aurora B activity and long abscission in naïve ESCs are due to high Wnt signaling

### Microtubules are transiently stabilized in slowly abscising cells

Chiron removal led to decreased Aurora B activity, less bridges, and shorter microtubule bridges, suggesting a crosstalk between Aurora B, microtubules, and abscission. Furthermore, Aurora B can regulate microtubule dynamics (50, 51) and microtubules need to be severed for abscission. We therefore hypothesized that differences in Aurora B activity between naïve and exiting cells could control abscission dynamics through regulation of microtubules. To test this, we imaged microtubules in maturing bridges in naïve and exiting cells. As expected, since the bridge narrows over time (1), microtubule intensity in the bridge decreased over time until abscission (Figure 3A,B, Movie 1,3). Cells can maintain microtubules over time by stabilizing them (52). To test whether microtubules are more stable in naïve cells, we used markers for post-translational modifications that predominantly decorate stable or dynamic microtubules (52). We stained naïve and exiting ESCs at different stages of bridge maturation for total tubulin, acetylated tubulin, which marks stable microtubules, and tyrosinated microtubules, which marks dynamic microtubules. Bridges were very dense structures containing acetylated and tyrosinated tubulin (Figure 3C, Supplementary Figure 3A). However, whilst total tubulin and tyrosinated tubulin intensity decreased over time (Figure 3C,D, Supplementary Figure 3A,B), acetylated tubulin intensity increased before decreasing (Figure 3C,E). This increase was much more pronounced in naïve than in exiting ESCs. This suggests a previously undescribed stage of transient microtubule stabilization during bridge maturation, which is more pronounced in naïve cells and correlates with slow abscission. To confirm this finding, we used a recently developed marker (StableMARK) to visualize stable microtubules in live cells (53). Using StableMARK, we confirmed that the intensity of stable microtubules increases then decreases over time in naïve and exiting ESC (Figure 3F-K, Movie 5,6, Supplementary Figure 3C,D). Of note, StableMARK expressing naïve and exiting cells displayed a delayed abscission compared to non-expressing cells consistent with previous findings that StableMARK can induce microtubule stabilization at high expression levels (53) (Figure 3G, 11.6h +/-4.6h for naïve cells versus 5.5h +/-1.4h in exiting cells, compare to Figure 3B). Finally, we wondered whether this transient increase in stable microtubules at the bridge was a general feature of dividing cells. To test this, we imaged a previously established cell line stably expressing StableMARK (53) and found that contrary to ESCs, U2OS expressing low levels of StableMARK had stable microtubules at the point of contact between the bridge and the cell bodies of the daughter cells, but only few around the midbody throughout bridge maturation (Supplementary Movie 6, Supplementary Figure 3E,F). Thus, the transient stabilization of bridge microtubules along the length of the bridge is not conserved between cell lines and could be a specific feature of stem cells.

**Fig. 3:**
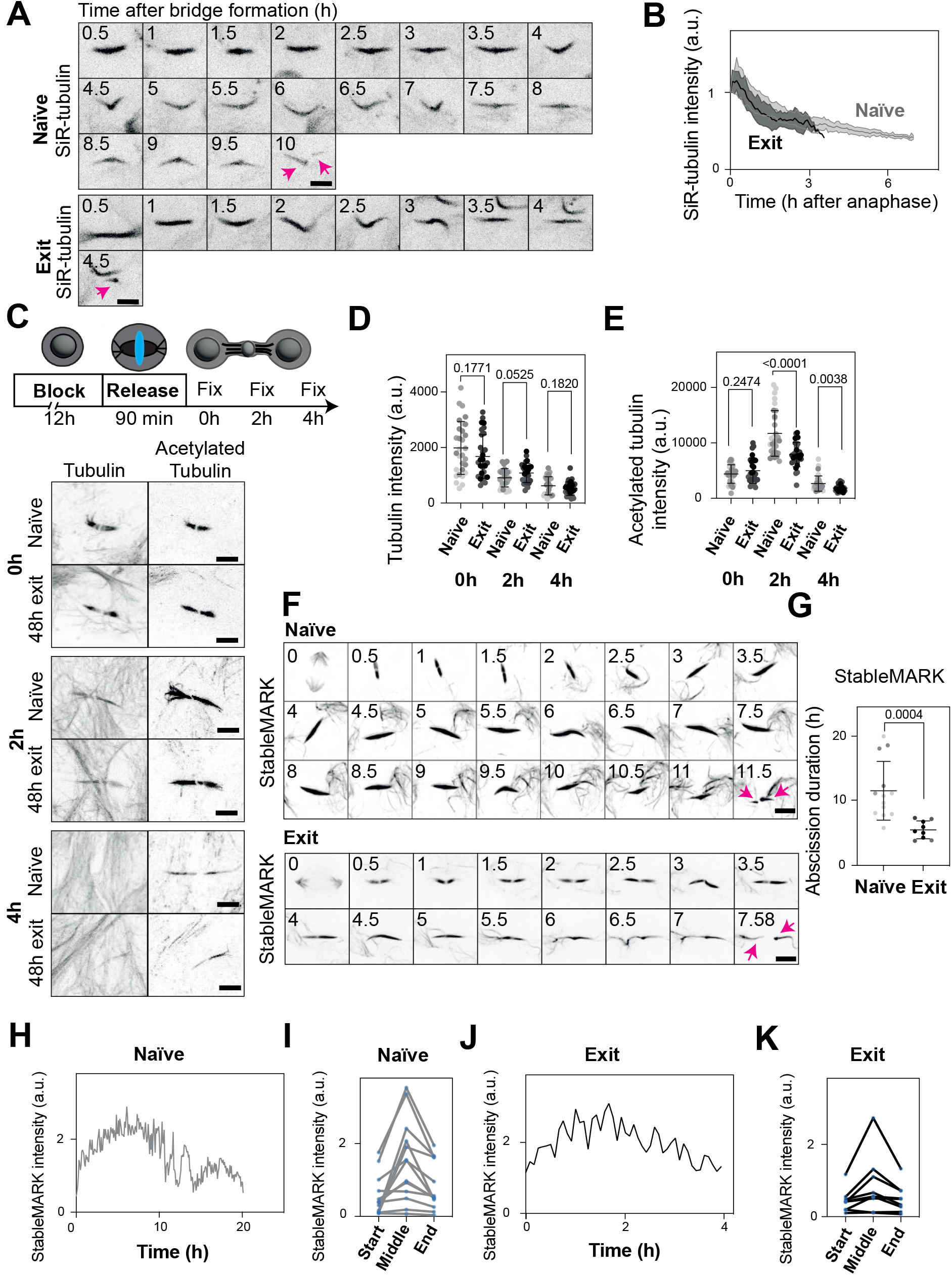
Microtubules are stabilized during slow abscission. A) Live-cell imaging of naïve (top) and 48h exiting (bottom) ESCs incubated with 20 nM SiR-tubulin. A Z-projection over the height of the whole cell is shown. One frame is shown every 30 min. The pink arrows indicate the cut sites (note that in the exiting example only one arm remain in frame after cut). The pink arrows indicate the cut sites. Scale bars: 5 *μ*m. B) Quantification of the intensity of SiR-tubulin over time. The mean and standard deviation are shown. N=3 replicates. C) Immunofluorescence showing the localization of total tubulin (left) and acetylated tubulin (middle) in naïve (top) and 48h exiting (bottom) ESCs during bridge maturation at bridge formation (0h), 2h after bridge formation (2h) and 4h after bridge formation (4h). The bridge is shown with the staining of total tubulin. A Z-projection over the height of the whole cell is shown. Scale bars: 5 *μ*m. D) Quantification of tubulin intensity in naïve (light grey) and exiting ESCs (dark grey) during bridge maturation. The mean and standard deviation are shown. N=3 replicates. E) Quantification of acetylated tubulin intensity in naïve (light grey) and exiting ESCs (dark grey) during bridge maturation. The mean and standard deviation are shown. N=3 replicates. F) Live-cell imaging of naïve (top) and 48h exit (bottom) ESCs transfected with StableMARK. A Z-projection over the height of the whole cell is shown. The pink arrows indicate the cut sites. One frame is shown every 30 min. Scale bars: 5 *μ*m. G) Graph showing the duration of abscission in naïve (light grey) or exiting (dark grey) ESCs expressing StableMARK. The mean and standard deviation are shown. N=3 replicates. H) Graph showing an example of the evolution of StableMARK intensity in a naïve ESC over time. The rest of the analyzed cells are shown in Supplementary Figure 3C. I) Graph showing the evolution of the intensity of StableMARK throughout bridge maturation of naïve ESCs, using bridge formation as “Start”, the last frame before bridge cutting as “End”, and the frame halfway these two points as “Middle”. N=5 replicates. J) Graph showing an example of the evolution of StableMARK intensity in a 48h exiting ESC over time. The rest of the analysed cells are shown in Supplementary Figure 3D. K) Graph showing the evolution of the intensity of StableMARK throughout bridge maturation in 48h exiting ESCs, using bridge formation as “Start”, the last frame before bridge cutting as “End”, and the frame halfway these two points as “Middle”. N=5 replicates.

### Aurora B activity controls microtubule stability

We then wondered whether Aurora B could control microtubule abundance and stability. To test this, we inhibited Aurora B activity with ZM and stained for total tubulin. We first noticed that bridges of cells treated with ZM were overall much shorter than control bridges (Figure 4A). Consistently, cells treated with ZM displayed less total tubulin, acetylated tubulin (Figure 4A-D), and tyrosinated tubulin (Supplementary Figure 4A-C), suggesting that inhibiting Aurora B activity decreases the amount of microtubules. We then decided to turn to live microscopy to test how the different populations of microtubules evolve over time. We analyzed the decrease in SiR-tubulin intensity (as a read-out for all microtubules) in naïve control cells and cells treated with ZM (Movie 1-2, Figure 1G,H). We found that, while the intensity of SiR-tubulin was overall lower in ZM-treated cells compared to control (Supplementary Figure 4, bottom panel) consistent with our fixed data, when we normalized each curve with the first time point set as 1, the intensity of SiR-Tubulin decreased over time with similar dynamics in ZM-treated naïve cells and in controls (Supplementary Figure 4D, top panel). Crucially, abscission always happened when microtubule intensity reached a similar threshold (Supplementary Figure 4D, bottom). We then imaged specifically stable microtubules with StableMARK in naïve ESC, and found that inhibiting Aurora B activity had a strong effect on stable microtubules, as the intensity of StableMARK decreased faster when cells were treated with ZM compared to control (Figure 4E-F, Movie 7,8). In particular, cells did not display the transient increase in stable microtubules described before (Figure 3F, H-K). Altogether, our data indicate that high Aurora B activity leads to an increase in the abundance of microtubules, in particular the stable population.

**Fig. 4:**
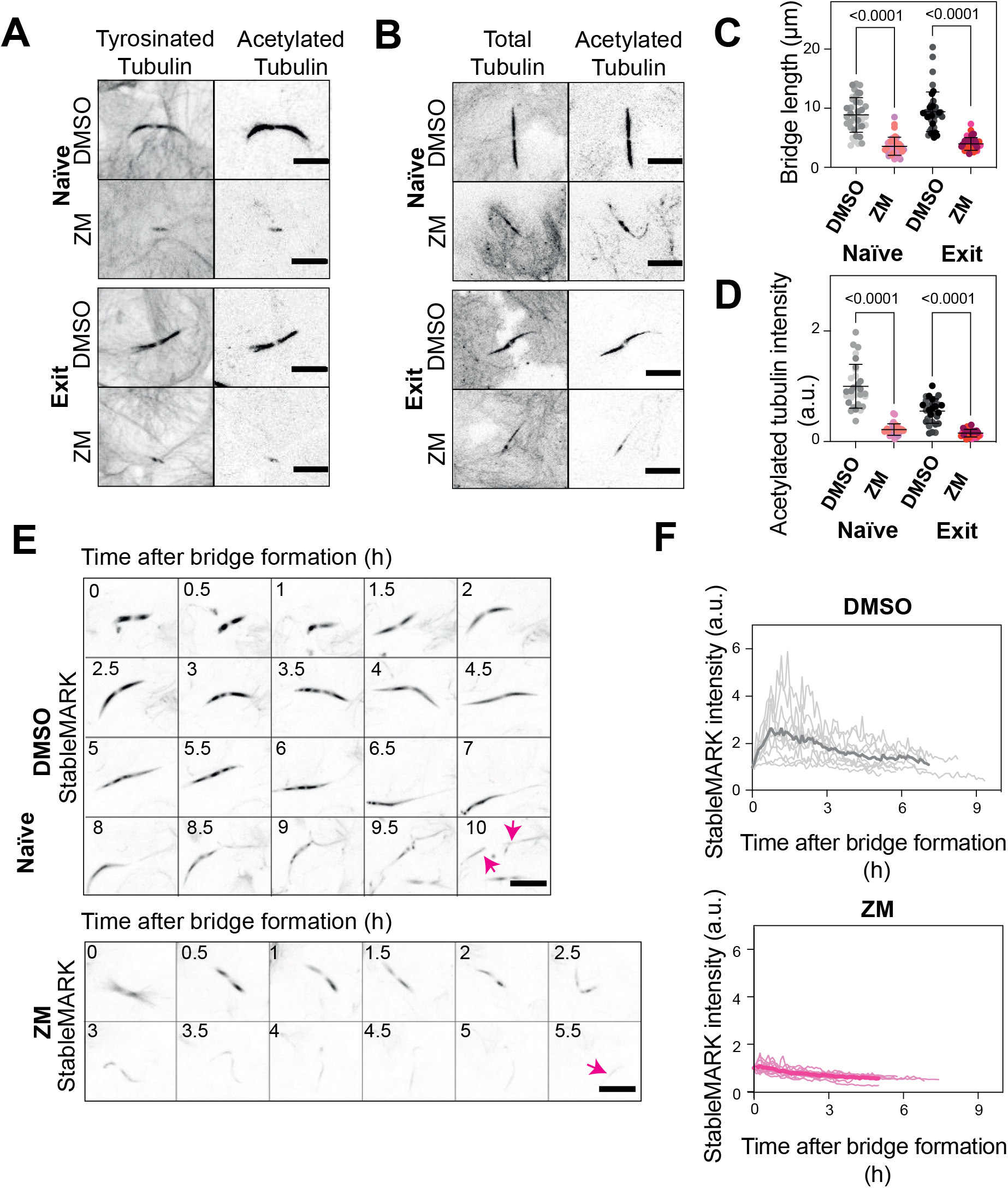
Aurora B activity controls microtubule stability. A) Immunofluorescence showing the size of naïve (top) or 48h exiting (bottom) bridges of ESCs treated for 90 min with DMSO or 2 *μ*M ZM447439. Bridges are stained for tyrosinated (left) or acetylated (right) tubulin. A Z-projection over the height of the whole cell is shown. Scale bars: 5 *μ*m. B) Immunofluorescence showing total tubulin (left) and acetylated tubulin (middle) in naïve (top) or 48h exiting (bottom) bridges of ESCs treated for 90 min with DMSO or 2 *μ*M ZM447439. A Z-projection over the height of the whole cell is shown. Scale bars: 5 *μ*m. C) Quantification of the length of bridges in naïve or 48h exiting ESCs treated with DMSO (light grey and dark grey respectively) or 2 *μ*M ZM447439 for 90 min (light pink and dark pink respectively). The mean and standard deviation are shown. N=3 replicates. D) Quantification of the intensity of acetylated tubulin in naïve or 48h exiting ESCs treated with DMSO (light grey and dark grey respectively) or 2 *μ*M ZM447439 for 90 min (light pink and dark pink respectively). The mean and standard deviation are shown. N=3 replicates. E) Live-cell imaging of naïve (top) and 48h exiting (bottom) ESCs expressing StableMARK. A Z-projection over the height of the whole cell is shown. Tubulin is shown in black. The pink arrows indicate the cut sites (note that in the ZM treated example only one arm remain in frame after cut). One frame is shown every 30 min. Scale bars: 5 *μ*m. F) Quantification of the intensity of StableMARK over time in naïve ESC expressing StableMARK and treated with DMSO (top, shades of grey and mean in darker grey) or 2 *μ*M ZM447439 for 90 min (bottom, shades of pink and mean in darker pink). The mean and individual curves are shown. N=3 replicates.

### Microtubule stability controls abscission speed

Since Aurora B controls both microtubule stability and abscission speed, we hypothesized that microtubule stability could control abscission speed. To test the role of stable microtubules in abscission speed, we first stabilized microtubules using Taxol. Stabilizing microtubules led to a small increase in Aurora B intensity in exiting ESCs only (Supplementary Figure 5A,B) and to a strong increase in P-Aurora B intensity in both naïve and exiting ESCs (Figure 5A,B), suggesting that stabilizing microtubules leads to an increase in Aurora B activity. Stabilizing microtubules also leads to an increase in the number of bridges (Figure 5C,D), suggesting that stabilizing microtubules slows abscission. To confirm this, we treated 48h exiting ESCs expressing Tubulin-GFP with Taxol and imaged cells during bridge maturation. Whilst control cells divided normally, abscission was severely delayed or never happened in cells treated with Taxol (Supplementary Movie 7,8). These data suggest that microtubule stability prevents abscission. To confirm that microtubule stability, not microtubule amount, controls abscission speed, we treated cells with Nocodazole, a drug that depolymerizes microtubules. Naïve ESCs were treated with increasing doses of Nocodazole. At low doses of nocodazole (0.5 *μ*M and 1 *μ*M), the acetylated microtubules largely remained intact, while there was a decrease in tyrosinated microtubules (Supplementary Figure 5C-G). At this concentration, the number of bridges was not affected (Figure 5E,F). At higher doses of Nocodazole, we noticed a decrease in the amount of acetylated microtubules (Supplementary Figure 5C-G), as well as a decrease in the number of bridges per cell (Figure 5E,F). Consistently with our data on Taxol-stabilized bridges, we also saw a decrease in P-Aurora B amount only when the number of acetylated microtubules decreased (Supplementary Figure 5G). Altogether, our data show that stable microtubules prevent abscission.

**Fig. 5:**
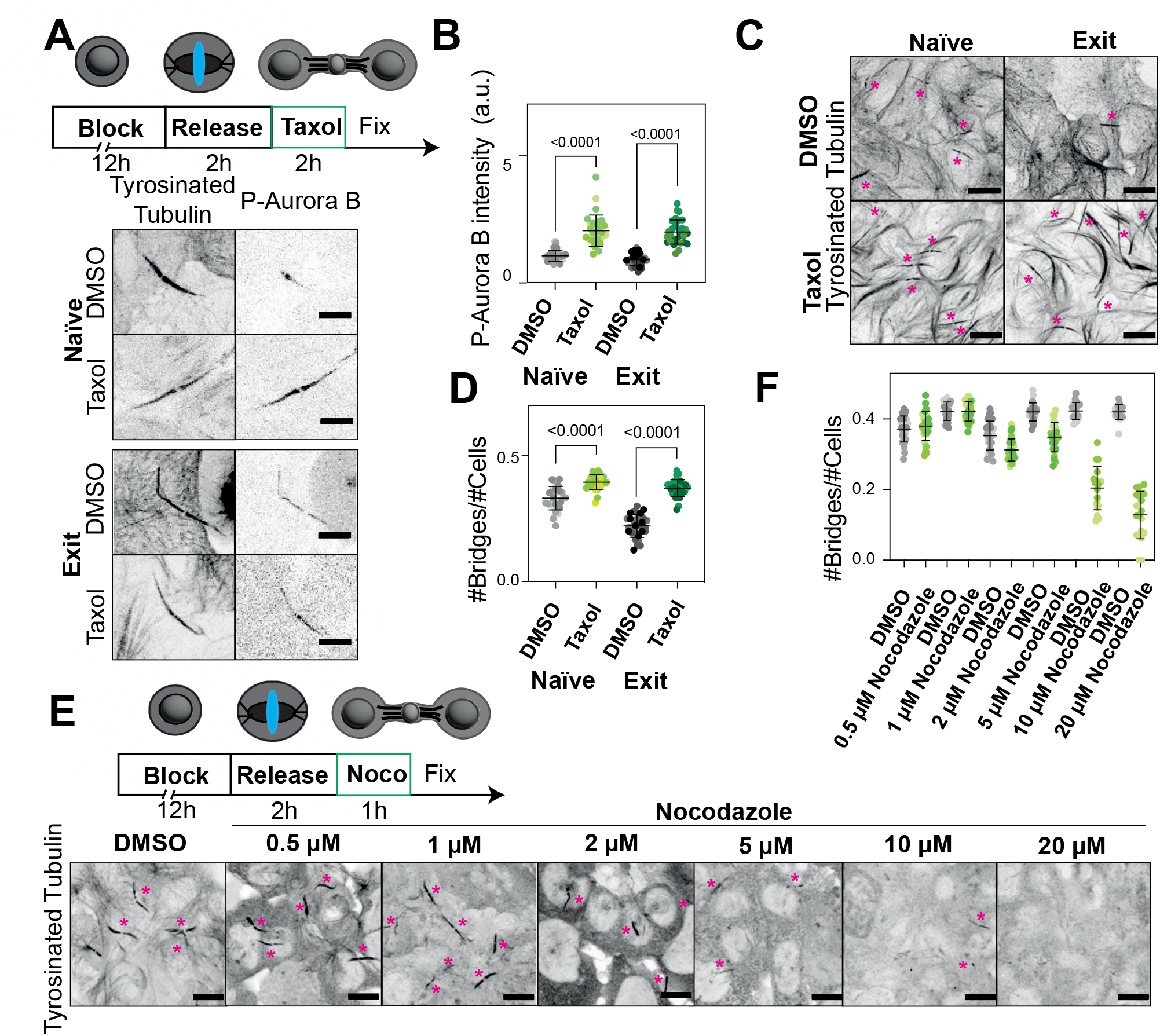
Microtubule stability delays abscission. A) Immunofluorescence showing the localization of P-Aurora B in naïve (top) and 48h exiting ESCs (bottom) treated with DMSO or 0.5 *μ*M Taxol. The bridge is shown with the staining of tyrosinated tubulin. A Z-projection over the height of the whole cell is shown. Scale bars: 5 *μ*m. B) Quantification of P-Aurora B intensity in naïve ESCs treated with DMSO or 0.5 *μ*M Taxol (light grey and light green, respectively) and exiting ESCs treated with DMSO or 0.5 *μ*M Taxol (dark grey and dark green, respectively). The mean and standard deviation are shown. N=3 replicates. C) Immunofluorescence showing the number of bridges in naïve (left) or 48h exiting (right) ESCs treated with DMSO (top) or 0.5 *μ*M Taxol (bottom). A Z-projection over the height of the whole cell is shown. The bridges are shown with the staining of tyrosinated tubulin. Bridges are highlighted with pink asterisks. Scale bars: 10 *μ*m. D) Quantification of the number of bridges per cell in naïve ESCs treated with DMSO or 0.5 *μ*M Taxol (light grey and light green respectively) or 48h exiting ESCs treated with DMSO or 0.5 *μ*M Taxol (dark grey and dark green respectively). The mean and standard deviation are shown. N=3 replicates. E) Immunofluorescence showing the number of bridges in naïve ESCs treated with DMSO or various concentrations of Nocodazole. A Z-projection over the height of the whole cell is shown. The bridges are shown with the staining of tyrosinated tubulin. Bridges are highlighted with pink asterisks. Scale bars: 10 *μ*m. F) Quantification of the number of bridges per cell in naïve ESCs treated with DMSO (light grey) or various concentrations of Nocodazole (light green). The mean and standard deviation are shown. N=3 replicates.

### Microtubule abscission occurs at zones of low stability

Finally, to confirm the relationship between low microtubule stability and abscission, we imaged StableMARK on both arms of the cytoplasmic bridges and compared the intensity between arms. We noticed that abscission happened asymmetrically, consistent with previous reports of asymmetric bridge remnants in stem cells (Supplementary Movie 1) (54). Strikingly, shortly before and after abscission, the cut arm always displayed a lower intensity of StableMARK (Figure 6A-D). These data show that lower levels of stable microtubule directly and locally correlate with abscission. We thus propose that in stem cells, high Aurora B activity leads to stabilization of microtubules in the bridge, which prevent abscission. Locally, microtubules can be destabilized, which triggers asymmetric abscission on one arm only. To confirm this model, we fixed cells several hours after bridge formation (6h after release in exiting ESCs, 7h after release in naïve ESCs) and investigated whether we could find an asymmetry in P-Aurora B and acetylated tubulin intensity in old bridges close to abscission. Strikingly, we found bridges that had asymmetric arms with one arm showing a higher intensity of acetylated tubulin and P-Aurora B but no difference in tyrosinated tubulin intensity (Figure 6F). The asymmetry in arms was found for 31.3% of naïve cells and 33.8% of exiting cells. This asymmetry was not obvious at earlier stages of bridge maturation (Figure 1B, 3C). Thus, our data suggest that close to abscission, the activity of Aurora B decreases locally and triggers a local decrease in microtubule stability which allows abscission. Finally, we compared abscission dynamics of naïve ESCs across our different conditions: while inhibiting Aurora B activity speeds up abscission, expressing StableMARK, which can lead to microtubule stabilization (53), increased abscission duration; expressing StableMARK in cells with a decreased Aurora B activity led to a return to a slow abscission (Figure 6G). Altogether, our data show that the stability of microtubules controls Aurora B activity and abscission dynamics.

**Fig. 6:**
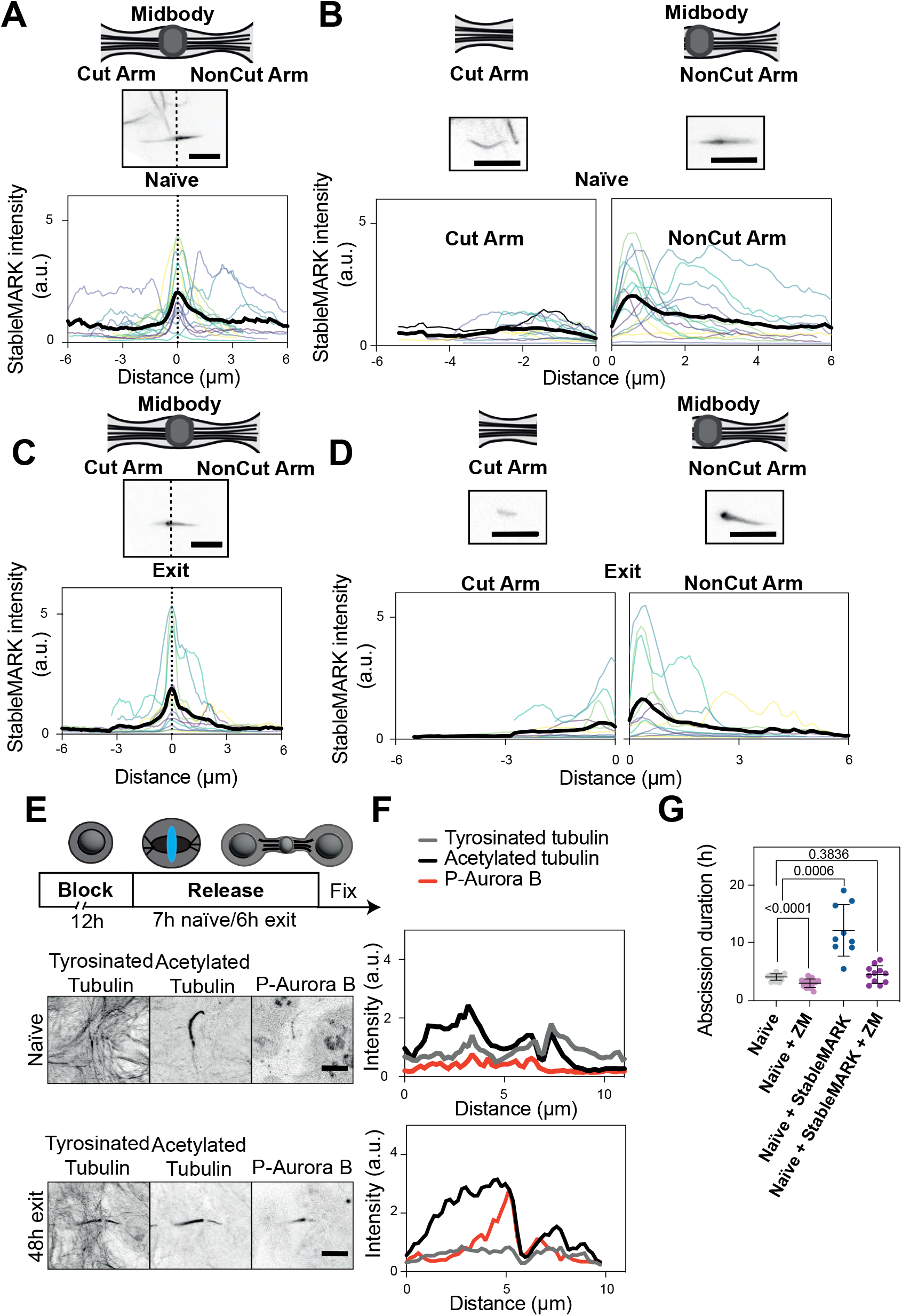
Bridges break at point of low microtubule stability. A) Top: cartoon showing a cytoplasmic bridge with the midbody in the center and the cut arm and non-cut arm before abscission. Bottom: graph showing the intensity of StableMARK along the two arms of bridges of naïve ESCs before abscission, with the cut arm on the left and the non-cut arm on the right. Insets show examples. Scale bars: 5 *μ*m. N=3 replicates. B) Top: cartoon showing a cytoplasmic bridge with the midbody in the center and the cut arm and non-cut arm after abscission. Bottom: graph showing the intensity of StableMARK along the two arms of bridges of naïve ESCs after abscission, with the cut arm on the left and the non-cut arm on the right. Insets show examples. Scale bars: 5 *μ*m. N=3 replicates. C) Top: cartoon showing a cytoplasmic bridge with the midbody in the center and the cut arm and non-cut arm before abscission. Bottom: graph showing the intensity of StableMARK along the two arms of bridges of 48h exiting ESCs before abscission, with the cut arm on the left and the non-cut arm on the right. Insets show examples. Scale bars: 5 *μ*m. N=3 replicates. D) Top: cartoon showing a cytoplasmic bridge with the midbody in the center and the cut arm and non-cut arm after abscission. Bottom: graph showing the intensity of StableMARK along the two arms of bridges of 48h exiting ESCs after abscission, with the cut arm on the left and the non-cut arm on the right. Insets show examples. Scale bars: 5 *μ*m. N=3 replicates. E) Immunofluorescence showing the localization of tyrosinated tubulin (left), acetylated tubulin (middle), and P-Aurora B (right) in naïve (top) and 48h exiting ESCs (bottom) 7h after release (naïve) or 6h after release (exit). The bridge is shown with the staining of tyrosinated tubulin. A Z-projection over the height of the whole cell is shown. Scale bars: 5 *μ*m. F) Line plot showing the intensity of tyrosinated tubulin, acetylated tubulin, and P-Aurora B in the two examples from panel E. Representative example from N=3 replicates. G) Graph showing the duration of abscission in naïve ESCs in several conditions: untreated and treated with 2 *μ*M ZM447439 for 90 min (similar to Figure 1H), expressing StableMARK (from Figure 3G) and expressing StableMARK and treated with 2 *μ*M ZM447439 for 90 min (from Figure 4F).

## Discussion

In this paper, we investigated the molecular mechanisms for the regulation of abscission dynamics in a physiological developmental context. We used mouse embryonic stem cells, which transition from a slow abscission to a fast abscission during exit from naïve pluripotency, to ask how the lifetime of cytoplasmic bridges is regulated.

Our data show that, apart from having a vital role in stalling abscission upon chromosome segregation errors (23, 38, 41, 43, 55–59), Aurora B kinase also regulates the dynamics of abscission during exit from naïve pluripotency. This is consistent with Aurora B activity maintaining cytoplasmic bridges in the early mouse embryo (40), or in human cells even in the absence of mitotic defects (43). Using precise temporal dissection of bridge maturation with fixed and live cell imaging, we showed that Aurora B activity decreases as the bridge matures and is higher in naïve cells compared to exiting cells (Figure 1). Inhibiting Aurora B activity drastically accelerates abscission (Figure 2). Altogether, our data show that increased Aurora B activity at the bridge slows down abscission in naïve embryonic stem cells.

What could explain the increased Aurora B activity at the bridge in naïve cells? Our data suggest a dual model, whereby naïve cells have higher amounts of Aurora B at the bridge as well as an increased activity of Aurora B. Indeed, the amount of Aurora B on the bridge is higher in naïve cells compared to exiting cells. At anaphase, Aurora B and its binding partners from the CPC are transported from the kinetochore along the microtubules (60) by the microtubule motor KIF20A (61). Our data suggest an increased transport or stability of Aurora B on the cytoplasmic bridge in naïve cells. Several hypotheses could explain this result. First, there is a global decrease in the expression levels of KIF20A during exit from naïve pluripotency (45, 62) which could explain the enhanced transport of Aurora B to the bridge in naïve cells. Moreover, the amount of microtubules present in the bridge can directly impact the abundance and activity of Aurora B (63); the increased stability of microtubules in naïve cells could promote the continuous recruitment and activation of Aurora B. Second, we show that the Wnt signaling pathway directly affects the abundance and activity of Aurora B. Indeed, dampening the Wnt pathway which is active in naïve cells (64) led to accelerated abscission and a reduction in Aurora B amount and activity. How the Wnt pathway controls Aurora B is unknown, but GSK3 (which is the protein targeted by Chiron) is known to phosphorylate and inhibit Aurora A (65), suggesting that it could similarly regulate Aurora B, although data in cancer cells indicate that different GSK3 inhibitors have no effect on Aurora B levels or phosphorylation during metaphase (66). In cancer cells, it has been proposed that Aurora B is degraded at the bridge to promote abscission (8), and Wnt signaling has been proposed to prevent protein degradation (67). Thus, high Wnt signaling in naïve cells could promote Aurora B stability. Finally, Wnt signaling can also locally stabilize microtubules (68, 69), suggesting that high Wnt signaling in naïve cells could participate to slow abscission through direct stabilization of bridge micro-tubules. Interestingly, dampening of Wnt signaling promotes binucleation in hepatocyte organoids (49), indicating that the Wnt signaling could play a conserved role in the regulation of cytokinesis.

We demonstrate that Aurora B plays a role in maintaining the naïve cell state (Figure 3). Indeed, a short burst of inhibition of Aurora B during exit from naïve pluripotency accelerates the fate transition, while OKA treatment (causing a decrease in Aurora B activity) slows exit down. We have previously shown that abscission speed is tightly coupled to cell fate and that the presence of the midbody slows down exit from naïve pluripotency (1). Thus, Aurora B could play an indirect role in cell fate by regulating abscission; however, our data do not exclude a model in which Aurora B also directly controls cell fate, and further studies will be necessary to untangle the two possibilities.

Our data show that Aurora B regulates the duration of abscission by maintaining microtubule stability (Figure 3-4). Indeed, we discover a previously undescribed transient increase in microtubule stability during maturation that depends on Aurora B activity. We do not know how Aurora B activity controls microtubule dynamics, but several hypotheses can be proposed. First, Aurora B or more broadly the CPC could regulate enzymes that modify tubulin post-translationally such as microtubule acetylases, for example, *α*-TAT1 (70, 71) (the acetyltransferase responsible for microtubule acetylation at Lysine 40), or microtubules deacetylases HDAC-6 (72) or SIRT2 (73). Second, Aurora B inactivates the minus end microtubule depolymerase MCAK by phosphorylation (50, 74). Thus, Aurora B could promote microtubule stability by inhibiting MCAK-dependent microtubule depolymerization. Interestingly, while lower Aurora B activity leads to a transient increase in the number of stable microtubules, it also leads to a global decrease in the number of microtubules. This suggests that two mechanisms exist, one where the activity of Aurora B controls microtubule abundance (perhaps through MCAK) and one where the activity of Aurora B controls microtubule stability ((consistent with the fact that MCAK prefers tyrosinated microtubules (75)).

We demonstrate that stable microtubules inhibit abscission (Figure 5-6). Indeed, increasing microtubule stability with Taxol led to slower abscission. Moreover, stabilizing microtubules with the expression of higher levels of StableMARK also slowed abscission. It is important to point out that while StableMARK could stabilize microtubules at high expression levels, in agreement with previous findings (53), we could still notice differences in abscission dynamics between naïve and exiting cells. Finally, depolymerizing dynamic microtubules with low doses of Nocodazole did not impair abscission dynamics, but higher doses of Nocodazole that perturb stable microtubules accelerated abscission. How could stable microtubules specifically delay abscission? We propose that microtubules prevent the primary constriction of the bridge. Consistent with this data, a recent preprint demonstrated that local severing of microtubules by a transient pool of actin polymerization is important for a timely abscission (**?** ). Stable microtubules are less sensitive to mechanical stress (76), thus stable microtubules could promote the structural integrity of the bridge. Interestingly, stable microtubules (which we demonstrate hinder abscission) are generally polyglutamylated (77) and polyglutamylation modulates spastin activity (78). In sum, our data suggest that abscission dynamics could depend on the regulation of spastin activity by microtubule stability.

Overall, our results demonstrate that the activity of Aurora B kinase can regulate abscission dynamics through the regulation of microtubule stability. Stable microtubules in turn promote Aurora B activity. In contrast to traditional models wherein microtubules are passive remnants of the mitotic spindle, our study identifies microtubules themselves as a main driver in abscission dynamics.

## ACKNOWLEDGEMENTS

We would like to thank all the members of the Chaigne lab and the whole department of Cell Biology, Neurobiology and Biophysics at Utrecht University, in particular Anna Akhmanova for helpful discussions. We also would like to thank Gautam Dey (EMBL), Susanne Lens (UCMU), Marie-Emilie Terret (Collège de France), and Ruby Peters (Cambridge University) for feedback on the manuscript.

## AUTHOR CONTRIBUTIONS

SK performed most of the experiments and analysis. AO performed some of the StableMARK experiments and analysis. EvB was involved in the experiments and analysis of the characterization of the different populations of microtubules in fixed samples. MI was involved in the imaging and analysis of fixed and live microtubules. WN was involved in the imaging and analysis of Aurora B and P-Aurora B. LK was involved in the conceptualization. AC performed some of the qPCRs and clonogenicity assays, and the conceptualization and supervision. All authors discussed the data and wrote the manuscript.

## FUNDING

This work was supported by Utrecht University and NWO (NWO-Vidi to AC).

## METHODS

### Data and materials availability

All data and materials used in the analysis are available upon request.

### Cell culture and transfection

#### Cell lines

E14 Mouse embryonic stem cells (a kind gift from Niels Geijsen, Hubrecht Institute) were used throughout the study. Rex1-GFP Gap43-mCherry (a kind gift from Kevin Chalut, Altos Cambridge) were used for Supplementary Movie 1, and U2OS stably expressing StableMARK (53) were used for Supplementary Figure 3 E,F. *Culture* E14 Mouse embryonic stem cells were routinely cultured on 10 cm cell culture dishes (Greiner Bio-one, 664160) coated with 0.1% gelatin/PBS in N2B27+2i-LIF with penicillin and streptomycin, at a controlled density (15-30k cells/cm square) and passaged every other day using Accutase (Sigma-Aldrich, #A6964). They were kept in 37C incubators with 5% CO2. Cells were regularly tested for mycoplasma. To trigger exit from naïve pluripotency, cells were plated in N2B27 after passaging, For a typical experiment, cells were passaged and plated onto laminin for 48h in either N2B27+2i-Lif (naïve) or N2B27 (exiting). The culture medium was made in house as described in (46) using DMEM/F-12, 1:1 mixture (Sigma-Aldrich, #D6421-6), Neurobasal medium (Life technologies #21103-049), 2.2 mM L-Glutamin (Thermofischer Scientific # 25030024), home-made N2 (see below), B27 (Life technologies #12587010), 3 μM Chiron (Merck #SML1046), 1 μM PD 0325901 (Merck #PZ0162), LIF (Merck # ESG1107), 0.1 mM –ethanol, 12.5 μg/mL Insulin zinc (ThermoFischer Scientific # 12585014). The 200 X home-made N2 was made using 8.791 mg/mL Apotransferrin (Sigma-Aldrich #T1147), 1.688 mg/mL Putrescine (Sigma-Aldrich #P5780), 3 μM Sodium Selenite (Sigma-Aldrich #S5261), 2.08 μg/mL Progesterone (Sigma-Aldrich #P8783), 8.8% BSA. For fixed and live imaging, the cells were plated on 35 mm round Ibidi dishes (IBI Scientific, #81156), coverslips in Invitrogen™ Attofluor™(#A7816) or 8-well Ibidi chambers (IBI Scientific #80807) coated with 10 mg/mL Laminin (Sigma #11243217001) overnight at 37C.

#### Transfection

For transfection, one well of an 8-well Ibidi chambers (IBI Scientific #80807) was coated with 10 mg/mL Laminin (Sigma, #11243217001) overnight at 37C. 1 μg of DNA was incubated in 50 μL Optimem (Thermofischer #11058021) for 5 min at room temperature. 1.2 μL of Lipofectamine™ (Thermofischer #18324012) was incubated at room temperature in 50 μL Optimem. The DNA and Lipofectamin solutions were combined and incubated at room temperature for 20 min. Meanwhile, the cells were passaged as described above and seeded in the well from which Laminin has been removed with 100 μL of culture media (N2B27+2i-Lif for Naïve cells or N2B27 for Exit cells). The transfection mix was added on top of the cells and the cells were placed in the incubator overnight, and typically rinsed the next morning and imaged in the afternoon.

### Drug treatment

#### Synchronization

When indicated, cells were synchronized using 10 μM RO-3306 for 15h and released for 90 min before further treatment. For the time course of bridge maturation, cells were fixed 2h after washing out the RO-3306 which corresponds to bridge formation (0h), or 2h or 4h later.

#### Inhibition of Aurora-B activity with ZM447439

When indicated, cells were treated with 2 μM ZM447439 (Selleckchem/BioConnect # S1103) or an equivalent amount of DMSO for 90 minutes, then rinsed 3 times. For live imaging ZM447439 was not rinsed.

#### Inhibition of Phosphatases PP1 and PP2A and B with Okadaic Acid

When indicated, cells were treated with 0.5 μM Okadaic Acid (Alomone lab #O-800) or an equivalent amount of DMSO for 60 minutes, then rinsed 3 times. For live imaging Okadaic Acid was not rinsed.

#### Stabilizing microtubules with Taxol

When indicated, cells were treated with 0,5 μM Taxol (Sigma-Aldrich # PHL89806) or an equivalent amount of DMSO for 1h, then fixed. For live imaging cells were treated with 0.5 μM Taxol at the beginning of imaging.

#### Depolymerization of microtubules with Nocodazole

When indicated, cells were treated with 0.5 μM, 1 μM, 2 μM, 5 μM, 10 μM or 20 μM Nocodazole or an equivalent amount of DMSO for 1h, then fixed.

### Immunofluorescence

The primary antibodies used were: alpha Tubulin Monoclonal Antibody (ThermoFischer Scientific #62204), P-Aurora B (ThermoFischer Scientific # PA5-105026), Aurora B AIM-1 (BD Biosciences # 611082), Citron Kinase (BD Biosciences #611376), Acetylated Tubulin (Merck # T7451 and Cell Signaling Technology # D20G3), Tyrosinated Tubulin (ThermoFischer # MA1-80017). The immunofluorescence was done as described in (79). Briefly, cells were fixed and permeabilised simultaneously using 4% Formaldehyde and 0.1% Triton in PBS for 10 minutes, rinsed 3 times in PBS then blocked in 3% BSA in PBS for 15 minutes. Primary antibodies were added at 1:200 in the blocking solution and incubated for 2h at room temperature. Cells were then rinsed 3 times in PBS then blocked for 15 minutes with the blocking solution and incubated with the secondary antibodies at 1:500 for 1h at room temperature. Cells were then rinsed 3 times with PBS, incubated for 10 minutes with Hoechst, rinsed 3 times with PBS, and kept in PBS until imaging.

For imaging of fixed samples a Zeiss LSM 519 700 confocal setup was used. It consists of an AxioObserver Z1 microscope with a Plan-Apochromat 520 63×/0.8 oil objective. The set-up was controlled using ZEN.

### Live imaging

For live imaging of microtubules, cells were incubated overnight with 20 nM SiR-tubulin (Spirochrome #SC002).

Live imaging was performed using a 60× (Plan Apo VC, NA 1.4; Nikon) oil-immersion objective on a Spinning Disc (Yokogawa CSU-X1-A1) Nikon Eclipse Ti microscope with Perfect Focus System equipped with a sample incubator (Tokai-Hit) and an Evolve 512 EMCCD camera (Photometrics), controlled with MetaMorph 7.7 software (Molecular Devices). Cobolt Calypso 491 nm and Cobolt 647 nm lasers were used for excitation. Images were acquired every 5 min.

### Colony-forming assay

Colony-forming assays were performed to test the dynamics of exit from naïve pluripotency. Cells were plated at low density (30k cells per well of a 24-well plate) onto plates coated with 0.1% gelatin in N2B27 for 20 hours. Cells were resuspended, counted, and plated at clonal density (300 cells per well of a 12-well plate) on 0.1% gelatin in N2B27+2i-LIF. After 5 days, the number of colonies was manually counted.

### qPCR

RNA extraction was performed using the RNA Easy Qiagen kit according to the manufacturer’s instructions. The reverse transcription was performed using the High-Capacity cDNA Reverse Transcription Kit (Thermofischer Scientific #4368814). qPCR was performed using SsoAdvancedTM Universal SYBR Green Supermix (BioRad, #172-5271), loading 2.3 mg per lane. Primers were bought from Integrated DNA technologies. The heatmap was generated using a web-based tool (80).

### Image analysis

All images were analysed in Fiji (National Institutes of Health, Bethesda, MD, USA) (81). Raw images were used for quantification. The contrast was adjusted for clarity of presentation in the figures.

#### Abscission duration scoring in live cells

Throughout the manuscript, we call abscission the moment where we see rupture of the whole lattice of microtubules in the bridge followed by a recoil of one or both side of the cut bridge. This is in line with our data showing that in ESC membrane rupture directly follow the rupture of the microtubule lattice (Supplementary Movie 1).

#### Number of bridges

To calculate the ratio between number of cells and number of bridges, the number of bridges was counted using the tubulin channel and the number of cells was counted using a marker of the nucleus (for example Hoechst or Aurora B). The number of bridges was then divided by the number of cells. A higher ratio suggests that abscission is slow.

#### Bridge length

To measure bridge length, a segmented line was used to trace the microtubules of the bridge from one cell to another cell. This measurement is used as bridge length.

#### Intensity of Aurora B/P-Aurora B/total tubulin / tyrosinated / acetylated tubulin

To measure intensities in intercellular bridges, a Region Of Interest (ROI) was created surrounding the bridge using one of the tubulin channel and the sum intensity projection was made of the slices in which a bridge was visible. Only flat bridges were selected. The mean Aurora B/P-Aurora B/tubulin intensity was measured using the ROI. To calculate the mean intensity of Aurora B/P-Aurora B/tubulin in the bridge, the background mean intensity was subtracted from the mean intensity of Aurora B/P-Aurora B/tubulin measured in the bridge.

#### Intensity of SiR-tubulin and StableMARK

To measure the intensity of SiR-tubulin and StableMARK in intercellular bridges over time, a maximum intensity projection was made of the slices in which a bridge was visible. A bleaching correction was applied to all images. This was done by using the Bleach Correction plugin in Fiji and the chosen correction method was Simple Ratio. The bridge was manually segmented in Fiji using the tubulin channel in every time frame and the SiRtubulin or StableMARK mean intensity was measured in that area. Only flat bridges were used. To calculate the mean intensity of SiR-tubulin and StableMARK in the bridge, the background mean intensity was subtracted from the mean intensity of SiR-tubulin/StableMARK measured in the bridge.

### Statistical analysis and data display

Data is given as mean±s.d., unless otherwise stated. Means of two groups were compared by an appropriate test (Student’s t-test if the data was normal with a similar standard deviation, with Welch correction if the standard deviation was different, or Mann-Whitney if the distribution was not normal). Anovas were perfomed in cases with more than 2 groups. In cases where representative intercellular bridge images are shown, similar observations were made in at least 10 bridges from at least 3 independent experiments. Different replicates are shown as variations of the main colors (for example, 3 shades of grey). Graphs were generated in GraphPad Prism (MathWorks, Natick, MA, USA). Fiji was used to scale images and adjust brightness and contrast. Figures were assembled in Adobe Illustrator (Adobe Systems, Mountain View, CA, USA).

## Notes

### Competing Interest Statement

The authors have declared no competing interest.

