## Supplementary Information for "Aurora B controls microtubule stability to regulate abscission dynamics in stem cells"

### Movie legends

**Movie 1.** Naïve ESC incubated with 20 nM SiR-tubulin and treated with DMSO during bridge maturation. A Z-projection over the height of the cell is shown. One frame is shown every 5 min. Scale bar: 10 $\mu$m.
**Movie 2.** Naïve ESC incubated with 20 nM SiR-tubulin and treated with 2 $\mu$M ZM447439 for 90 min during bridge maturation. A Z-projection over the height of the cell is shown. One frame is shown every 5 min. Scale bar: 10 $\mu$m.
**Movie 3.** 48h exiting ESC incubated with 20 nM SiR-tubulin and treated with DMSO during bridge maturation. A Z-projection over the height of the cell is shown. One frame is shown every 5 min. Scale bar: 10 $\mu$m.
**Movie 4.** 48h exiting ESC incubated with 20 nM SiR-tubulin and treated with 2 $\mu$M ZM447439 for 90 min during bridge maturation. A Z-projection over the height of the cell is shown. One frame is shown every 5 min. Scale bar: 10 $\mu$m.
**Movie 5.** Naïve ESC expressing StableMARK during bridge maturation. A Z-projection over the height of the cell is shown. One frame is shown every 5 min. Scale bar: 10 $\mu$m.
**Movie 6.** 48h exiting ESC expressing StableMARK during bridge maturation. A Z-projection over the height of the cell is shown. One frame is shown every 5 min. Scale bar: 10 $\mu$m.
**Movie 7.** Naïve ESC expressing StableMARK and treated with DMSO during bridge maturation. A Z-projection over the height of the cell is shown. One frame is shown every 5 min. Scale bar: 10 $\mu$m.
**Movie 8.** Naïve ESC expressing StableMARK and treated with 2 $\mu$M ZM447439 for 90 min during bridge maturation. A Z-projection over the height of the cell is shown. One frame is shown every 5 min. Scale bar: 10 $\mu$m.
**Supplementary Movie 1.** 48h exiting ESC stably expressing Rex1-GFP and Gap43-mCherry and incubated with 20 nM SiR-tubulin. One frame is shown every 5 min. The SiR-tubulin channel is shown on the left and the Gap43-mCherry on the right, showing that membrane rupture shortly follows microtubule rupture. Scale bar: 10 $\mu$m.
**Supplementary Movie 2.** Naïve ESC incubated with 20 nM SiR-tubulin and treated with DMSO during bridge maturation. A Z-projection over the height of the cell is shown. One frame is shown every 5 min. Scale bar: 10 $\mu$m.
**Supplementary Movie 3.** Naïve ESC incubated with 20 nM SiR-tubulin and treated with 0.5 $\mu$M Okadaic Acid during bridge maturation. A Z-projection over the height of the cell is shown. One frame is shown every 5 min. Scale bar: 10 $\mu$m.
**Supplementary Movie 4.** 48h exiting ESC incubated with 20 nM SiR-tubulin and treated with DMSO during bridge maturation. A Z-projection over the height of the cell is shown. One frame is shown every 5 min. Scale bar: 10 $\mu$m.
**Supplementary Movie 5.** 48h exiting ESC incubated with 20 nM SiR-tubulin and treated with 0.5 10 $\mu$M Okadaic Acid during bridge maturation. A Z-projection over the height of the cell is shown. One frame is shown every 5 min. Scale bar: 10 $\mu$m.
**Supplementary Movie 6.** U2OS cell stably expressing StableMARK during bridge maturation. A Z-projection over the height of the cell is shown. One frame is shown every 2.5 min. Scale bar: 10 $\mu$m.
**Supplementary Movie 7.** 48h exiting ESC expressing GFP-tubulin treated with DMSO during bridge maturation. A Z-projection over the height of the cell is shown. One frame is shown every 5 min. Scale bar: 10 10 $\mu$m.
**Supplementary Movie 8.** 48h exiting ESC expressing GFP-tubulin treated with 0.5 $\mu$M Taxol during bridge maturation. A Z-projection over the height of the cell is shown. One frame is shown every 5 min. Scale bar: 10 $\mu$m.

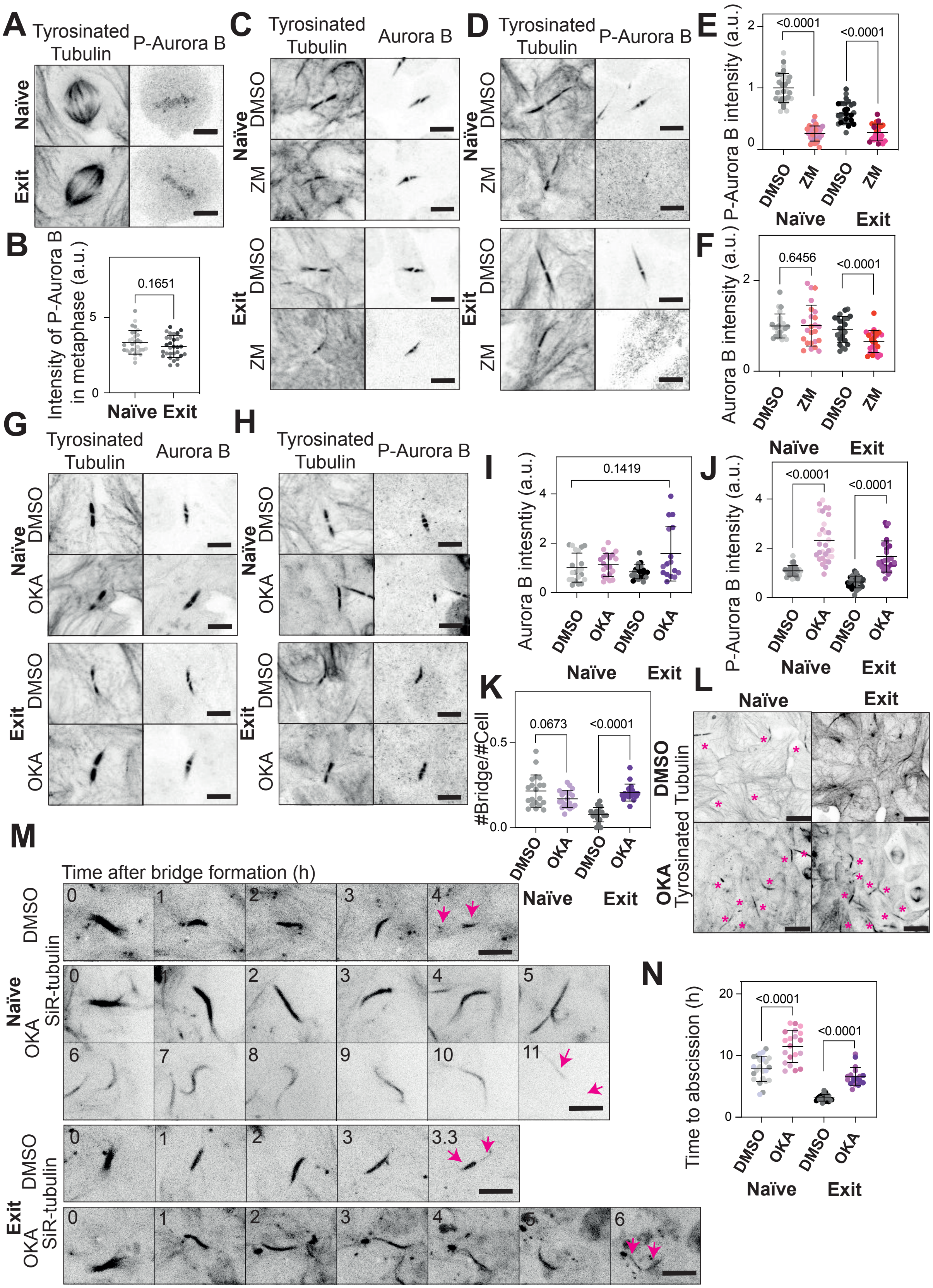


**Supplementary Figure 1. Phosphatase activity opposes Aurora B kinase in delaying abscission in ESC.** B) Immunofluorescence showing the localization of P-Aurora B in the metaphase plate of naïve (top) and 48h exiting ESCs (bottom). The bridge is shown with the staining of tyrosinated tubulin. A Z-projection over the height of the whole cell is shown. Scale bars: 5 $\mu$m. A) Quantification of P-Aurora B intensity in the metaphase plate of naïve ESC (light grey) and exiting ESCs (dark grey). The mean and standard deviation are shown. N=3 replicates. C) Immunofluorescence showing the localization of Aurora B in naïve (top) and 48h exiting ESCs (bottom) treated with DMSO or 2 $\mu$M ZM447439. The bridge is shown with the staining of tyrosinated tubulin. A Z-projection over the height of the whole cell is shown. Scale bars: 5 $\mu$m. D) Immunofluorescence showing the localization of P-Aurora B in naïve (top) and 48h exiting ESCs (bottom) treated with DMSO or 2 $\mu$M ZM447439. The bridge is shown with the staining of tyrosinated tubulin. A Z-projection over the height of the whole cell is shown. Scale bars: 5 $\mu$m. E) Quantification of Aurora B intensity in naïve ESC treated with DMSO or 2 $\mu$M ZM447439 (light grey and light pink, respectively) and exitsing ESCs treated with DMSO or 2 $\mu$M ZM447439 (dark grey and dark pink, respectively). The mean and standard deviation are shown. N=3 replicates. F) Quantification of P-Aurora B intensity in naïve ESC treated with DMSO or 2 $\mu$M ZM447439 (light grey and light pink, respectively) and exiting ESCs treated with DMSO or 2 $\mu$M ZM447439 (dark grey and dark pink, respectively). The mean and standard deviation are shown. N=3 replicates. G) Immunofluorescence showing the localization of Aurora B in naïve (top) and 48h exiting ESCs (bottom) treated with DMSO or 0.5 $\mu$M Okadaic Acid. The bridge is shown with the staining of tyrosinated tubulin. A Z-projection over the height of the whole cell is shown. Scale bars: 5 $\mu$m. H) Immunofluorescence showing the localization of P-Aurora B in naïve (top) and 48h exiting ESCs (bottom) treated with DMSO or 0.5 $\mu$M Okadaic Acid. The bridge is shown with the staining of tyrosinated tubulin. A Z-projection over the height of the whole cell is shown. Scale bars: 5 $\mu$m. I) Quantification of Aurora B intensity in naïve ESC treated with DMSO or 0.5 $\mu$M Okadaic Acid (light grey and light purple, respectively) and exiting ESCs treated with DMSO or 0.5 $\mu$M Okadaic Acid (dark grey and dark purple, respectively). The mean and standard deviation are shown. N=3 replicates. J) Quantification of P-Aurora B intensity in naïve ESC treated with DMSO or 0.5 $\mu$M Okadaic Acid (light grey and light purple, respectively) and exiting ESCs treated with DMSO or 0.5 $\mu$M Okadaic Acid (dark grey and dark purple, respectively). The mean and standard deviation are shown. N=3 replicates. K) Quantification of the number of bridges per cell in naïve ESC treated with DMSO or 0.5 $\mu$M Okadaic acid (light grey and light purple, respectively) and exiting ESCs treated with DMSO or 0.5 $\mu$M Okadaic acid (dark grey and dark purple, respectively). The mean and standard deviation are shown. N=3 replicates. L) Immunofluorescence showing the number of bridges in naïve (left) and 48h exiting ESCs (right) treated with DMSO (top) or 0.5 $\mu$M Okadaic Acid (bottom). A Z-projection over the height of the whole cell is shown. The bridges are shown with the staining of tyrosinated tubulin. Bridges are highlighted with pink asterisks. Scale bars: 10 $\mu$m. M) Live-cell imaging of naïve (top) and exiting (bottom) ESCs incubated overnight with 20 nM SiR-tubulin after addition of DMSO (top) or 0.5 $\mu$M Okadaic Acid (bottom). A Z-projection over the height of the whole cell is shown. Tubulin is shown in black. The pink arrows indicate the cut sites. One frame is shown every 60 min. Scale bars: 10 $\mu$m. N) Quantification of the duration of abscission from bridge formation until microtubule severing in naïve and exiting ESCs treated with DMSO or 0.5 $\mu$M Okadaic Acid. The mean and standard deviation are shown. N=3 replicates.

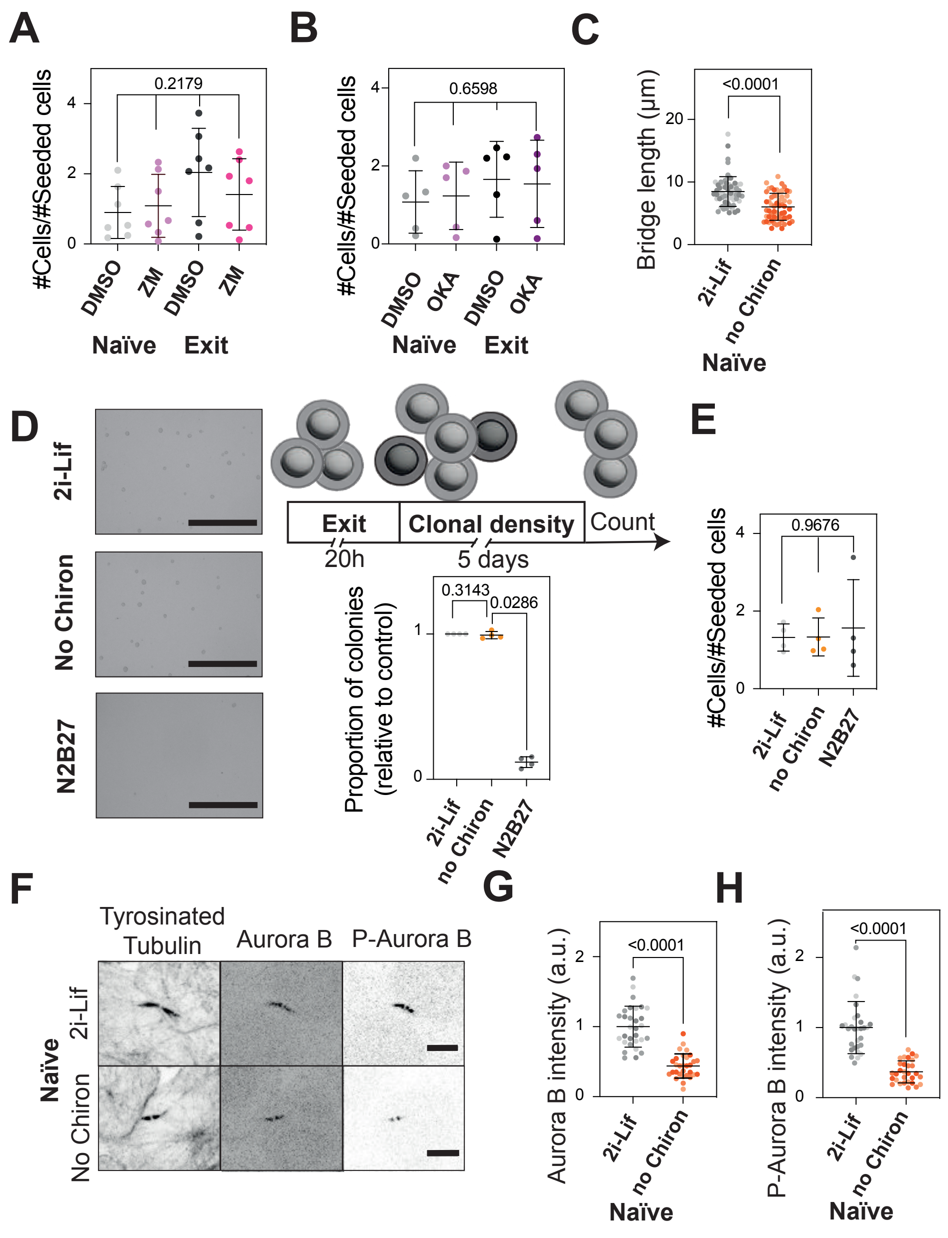


**Supplementary Figure 2. Wnt signaling promotes Aurora B activity and amount in the bridge.** A) Quantification of the proliferation rate in naïve ESC treated with DMSO or 2 $\mu$M ZM447439 (light grey and light pink, respectively) and exit ESC treated with DMSO or 2 $\mu$M ZM447439 (dark grey and pink, respectively). N= 4. B) Quantification of the proliferation rate in naïve ESC treated with DMSO or 0.5 $\mu$M Okadaic Acid (light grey and light purple, respectively) and exit ESC treated with DMSO or 2 $\mu$M ZM447439 (dark grey and purple, respectively). N= 4. C) Quantification of the bridge length in naïve ESC in normal 2i-Lif media or after Chiron withdrawal (light grey and orange, respectively). The mean and standard deviation are shown. N= 3. D) Clonogenicity assay on ESC in 2i-Lif, without Chiron, or in N2B27. Left panel: colonies formed by control cells after 20h in 2i-Lif, without Chiron or during exit from naive pluripotency (N2B27). Scale bars: 1 mm. Right: quantification of the number of colonies formed. N= 4. E) Quantification of the proliferation rate after 20h for naïve ESC with or without Chiron or exit cells (N2B27). N= 4 F) Immunofluorescence showing the localization of Aurora B and P-Aurora B in normal 2i-Lif media (top) and after Chiron withdrawal (bottom). The bridge is shown with the staining of tyrosinated tubulin. A Z-projection over the height of the whole cell is shown. Scale bars: 5 $\mu$m. G) Quantification of Aurora B intensity in normal 2i-Lif media and after Chiron withdrawal (light grey and orange, respectively). The mean and standard deviation are shown. N=3 replicates. H) Quantification of P-Aurora B intensity in normal 2i-Lif media and after Chiron withdrawal (light grey and orange, respectively). The mean and standard deviation are shown. N=3 replicates.

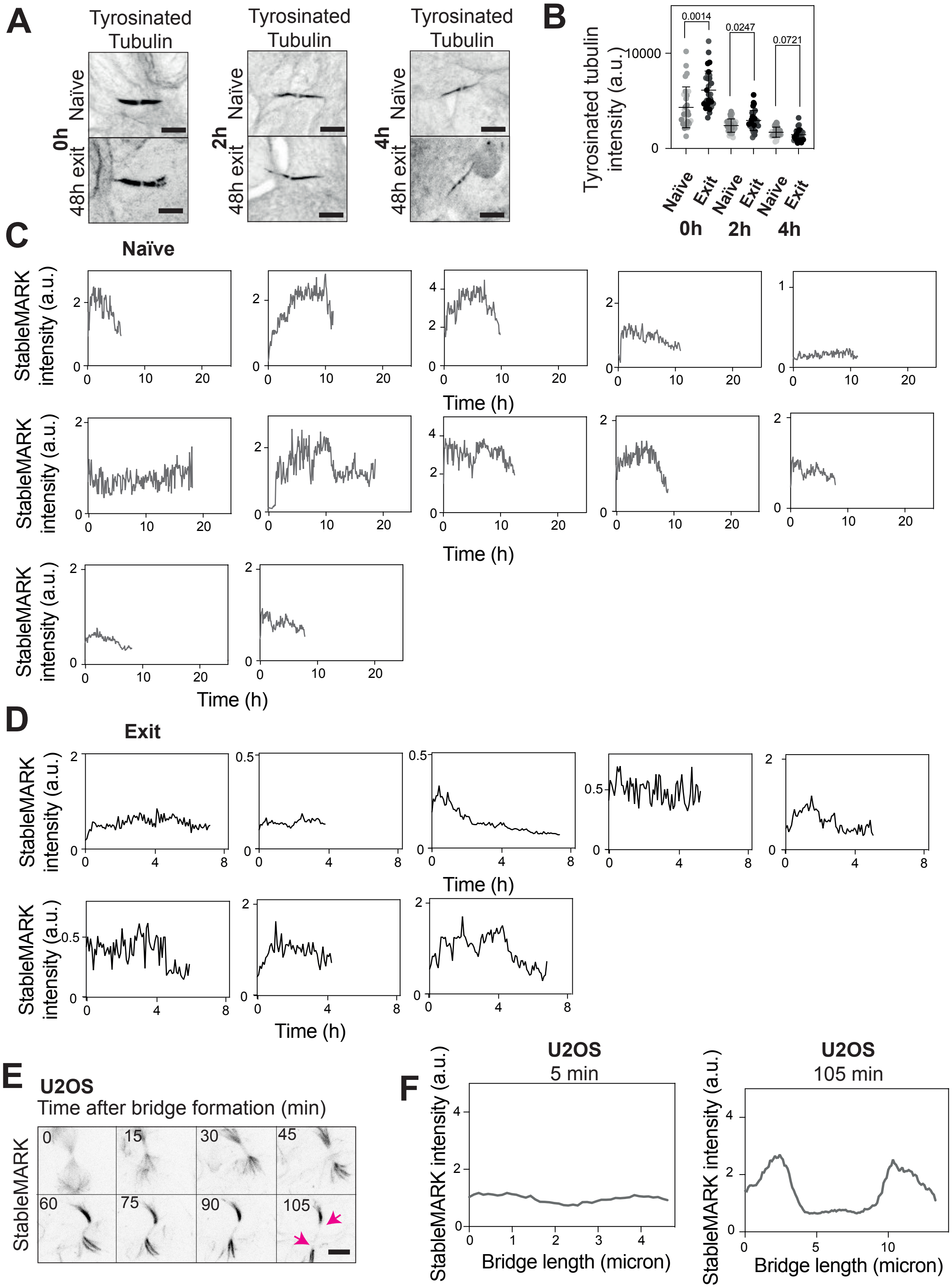


**Supplementary Figure 3. Changes in microtubule stability during bridge maturation.** A) Immunofluorescence showing the localization of tyrosinated tubulin in naïve (top) and 48h exit (bottom) ESC during bridge maturation at bridge formation (0h), 2h after bridge formation (2h) and 4h after bridge formation (4h). The bridge is shown with the staining of tyrosinated tubulin. A Z-projection over the height of the whole cell is shown. Scale bars: 5 $\mu$m. B) Quantification of tyrosinated tubulin intensity in naïve (light grey) and exit ESC (dark grey) during bridge maturation. The mean and standard deviation are shown. N=3 replicates. C) Quantification of the intensity of StableMARK over time in naïve ESC for single cells. N=12 replicates. D) Quantification of the intensity of StableMARK over time in exit ESC for single cells. N=8 replicates. E) Live-cell imaging of U2OS cells stably expressing StableMARK. A Z-projection over the height of the whole cell is shown. One frame is shown every 15 min. The pink arrows indicate the cut sites. Scale bars: 5 $\mu$m. F) Quantification of intensity of StableMARK along the bridge of the U2OS cell presented in E) 5 min (left panel) and 105 min (right panel) after bridge formation. Representative example of N=3 replicates.

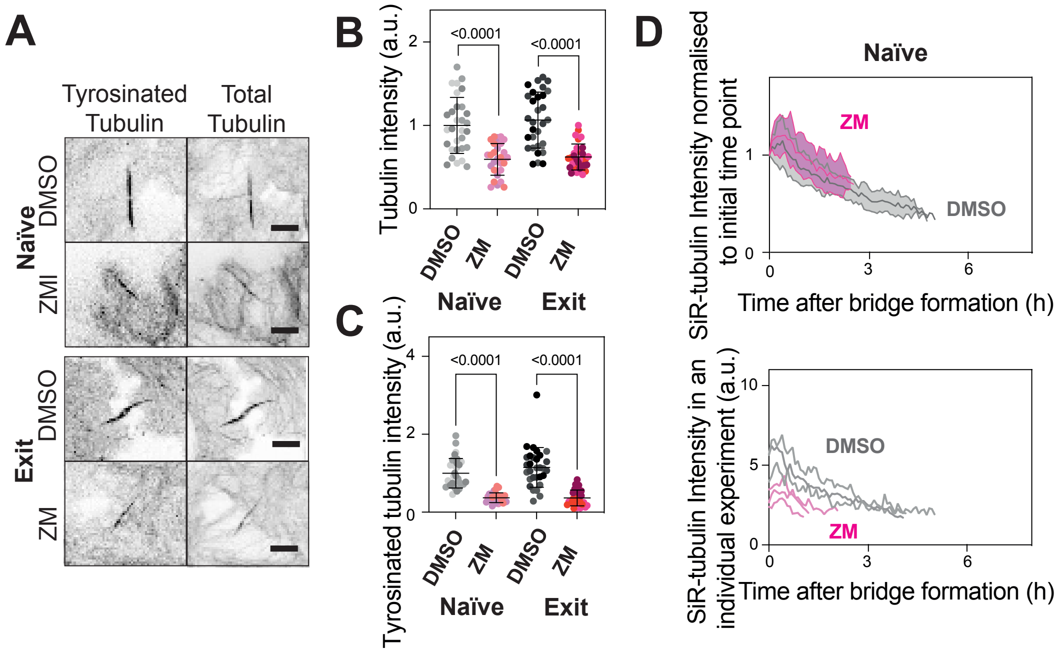


**Supplementary Figure 4. Aurora B influences microtubule amount and dynamics.** A) Immunofluorescence showing the localization of tyrosinated and total tubulin in the bridge in naïve (top) and 48h exit ESC (bottom) treated with DMSO or 2 $\mu$M ZM447439. A Z-projection over the height of the whole cell is shown. Scale bars: 5 $\mu$m. B) Quantification of total tubulin intensity in naïve ESC treated with DMSO or 2 $\mu$M ZM447439 (light grey and light red, respectively) and exit ESC treated with DMSO or 2 $\mu$M ZM447439 (dark grey and dark red, respectively). The mean and standard deviation are shown. N=3 replicates. C) Quantification of tyrosinated tubulin intensity in naïve ESC treated with DMSO or 2 $\mu$M ZM447439 (light grey and light red, respectively) and exit ESC treated with DMSO or 2 $\mu$M ZM447439 (dark grey and dark red, respectively). The mean and standard deviation are shown. N=3 replicates. D) Quantification of the intensity of SiR-tubulin over time in naïve ESC treated with DMSO or 2 $\mu$M ZM447439 (light grey and light red, respectively). The mean and standard deviation are shown, and each curve is normalized to the first time point which is set at 1. N=3 replicates. Bottom: individual curves for one replicate.

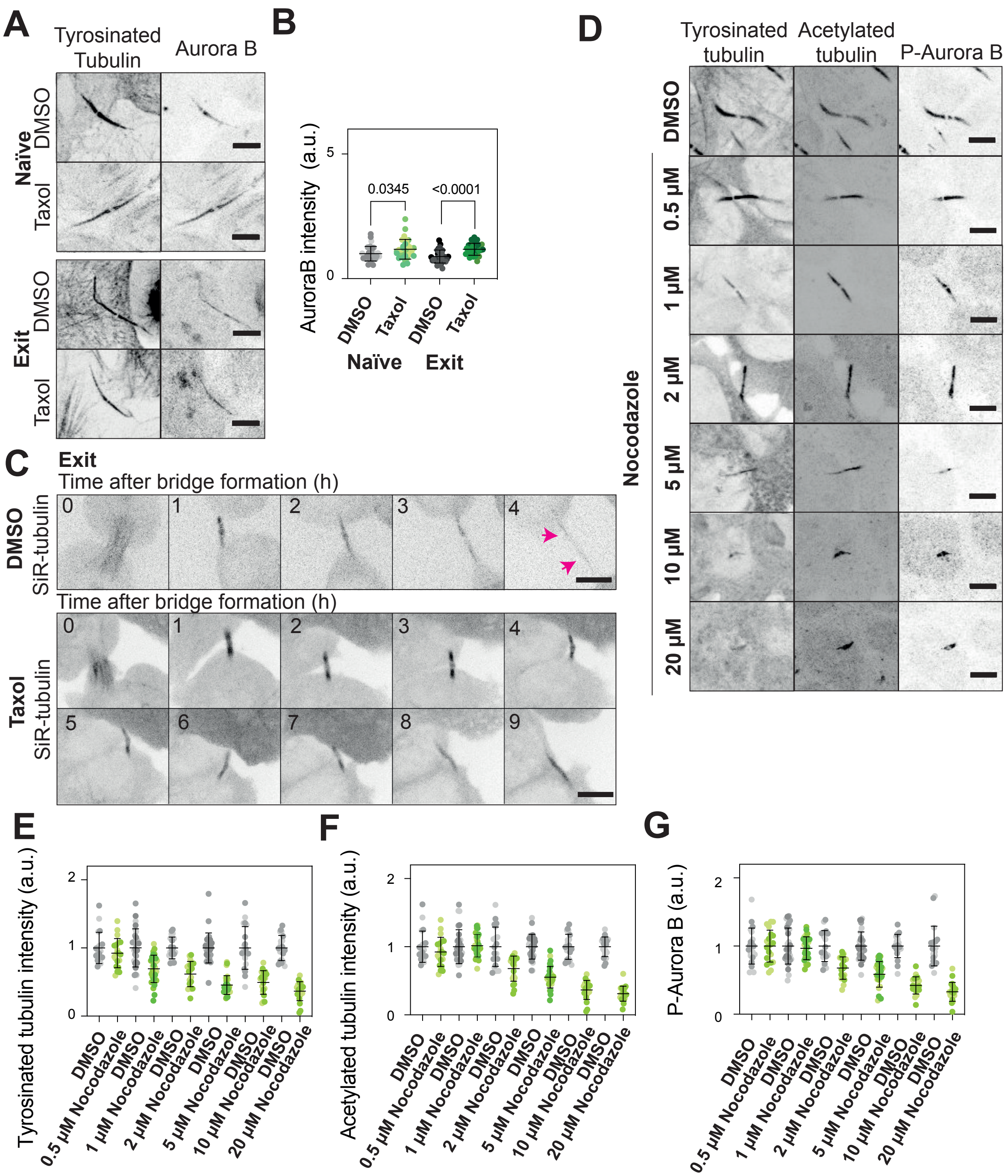


**Supplementary Figure 5. Microtubule stability and Aurora B act together in regulating abscission dynamics.** A) Immunofluorescence showing the localization of Aurora B in naïve (top) and 48h exit ESC (bottom) treated with DMSO or 0.5 $\mu$M Taxol. The bridge is shown with the staining of tyrosinated tubulin. A Z-projection over the height of the whole cell is shown. Scale bars: 5 $\mu$m. B) Quantification of Aurora B intensity in naïve ESC treated with DMSO or 0.5 $\mu$M Taxol (light grey and light green, respectively) and exit ESC treated with DMSO or 0.5 $\mu$M Taxol (dark grey and dark green, respectively). The mean and standard deviation are shown. N=3 replicates. C) Live-cell imaging of exit ESC transfected with GFP-tubulin after addition of DMSO (top) or 0.5 $\mu$M Taxol (bottom). A Z-projection over the height of the whole cell is shown. Tubulin is shown in black. One frame is shown every 60 min. The pink arrow indicates the cut site. Scale bars: 10 $\mu$m. D) Immunofluorescence showing the localization of tyrosinated tubulin, acetylated tubulin, and P-Aurora B in naïve ESC treated with DMSO or various concentrations of Nocodazole for 1h. A Z-projection over the height of the whole cell is shown. Scale bars: 5 $\mu$m. E) Quantification of acetylated tubulin intensity in naïve ESC treated with DMSO or various concentrations of Nocodazole (light grey and light green, respectively). The mean and standard deviation are shown. N=3 replicates. F) Quantification of tyrosinated tubulin intensity in naïve ESC treated with DMSO or various concentrations of Nocodazole (light grey and light green, respectively). The mean and standard deviation are shown. N=3 replicates. G) Quantification of P-Aurora B intensity in naïve ESC treated with DMSO or various concentrations of Nocodazole (light grey and light green, respectively). The mean and standard deviation are shown. N=3 replicates.
